# Unravelling the debate on heme effects in COVID-19 infections

**DOI:** 10.1101/2020.06.09.142125

**Authors:** Marie-Thérèse Hopp, Daniel Domingo-Fernández, Yojana Gadiya, Milena S. Detzel, Benjamin F. Schmalohr, Francèl Steinbock, Diana Imhof, Martin Hofmann-Apitius

## Abstract

The SARS-CoV-2 outbreak was recently declared a worldwide pandemic. Infection triggers the respiratory tract disease COVID-19, which is accompanied by serious changes of clinical biomarkers such as hemoglobin and interleukins. The same parameters are altered during hemolysis, which is characterized by an increase in labile heme. We present two approaches that aim at analyzing a potential link between available heme and COVID-19 pathogenesis. Four COVID-19 related proteins, i.e. the host cell proteins ACE2 and TMPRSS2 as well as the viral protein 7a and S protein, were identified as potential heme binders. We also performed a detailed analysis of the common pathways induced by heme and SARS-CoV-2 by superimposition of knowledge graphs covering heme biology and COVID-19 pathophysiology. Herein, focus was laid on inflammatory pathways, and distinct biomarkers as the linking elements. Finally, the results substantially improve our understanding of COVID-19 infections and disease progression of patients with different clinical backgrounds and expand the diagnostic and treatment options.

## 1 Introduction

In the beginning of 2020, the **co**rona**vi**rus **d**isease **19** (COVID-19) has been declared a pandemic of international concern and an unprecedented challenge for the country-specific health care systems(Cucinotta and Vanelli, 2020). COVID-19 is caused by infections with **s**evere **a**cute **r**espiratory **s**yndrome **co**rona**v**irus **2** (SARS-CoV-2) and is accompanied by pneumonia, acute respiratory distress syndrome (ARDS) associated with a cytokine storm, and death in the most severe cases (Ye et al., 2020; Zhou et al., 2020a). By taking a closer look into the molecular mechanisms of the infection and disease development, it is important to note that patients with severe COVID-19 often had a history of hypertension, yet also chronic kidney disease, cardiovascular disease or diabetes mellitus compared to those with milder disease progression (Ji et al., 2020; Zhou et al., 2020a). The scientific evidence of a potential higher risk for these patients, however, is still pending. Furthermore, there is evidence that the renin-angiotensin system (RAS), which is associated with hypertension, is directly associated with viral transmission (Hanff et al., 2020). An essential part of RAS is the enzyme angiotensin-converting enzyme 2 (ACE2) that is expressed on the cell surface of alveolar epithelial cells of the lungs (Ren, 2006). More precisely, a recent report identified specific bronchial transient secretory cells, a cell state between goblet (responsible for mucus production) and ciliated cells (responsible for airway clearance) in human bronchial epithelial cells to be primarily attacked during the viral infection (Lukassen et al., 2020; Xu et al., 2020).

The virus gains access to the host cell by docking of its spike proteins (S protein) to the membrane surface of the host cell, which primarily occurs via the transmembrane protein ACE2 (Hoffmann et al., 2020). This interaction between the host cell and SARS-CoV-related viruses is known since 2012 and involves the residues Gln394 of the S protein and Lys31 of ACE2 as the critical interacting amino acids (Wu et al., 2012). In this context, the membrane protein M (M protein) is discussed to be relevant for the entry and attachment of the virus by interacting with the S protein (Bianchi et al., 2020). Furthermore, the M protein may also be important for the budding process of the virus, since it interacts with the nucleocapsid envelope protein (E protein) and the S protein during virus particle assembly (Alsaadi and Jones, 2019). Additionally, it has been proposed that the E protein oligomerizes to form ion channels, and also plays a role in the assembly of the viral genome (Ruch and Machamer, 2012). Although protein 7a has not yet been fully characterized it is already known to act as an accessory protein (as derived by similarity from SARS-CoV), while interacting with M and E proteins (Fielding et al., 2006). This process is essential for virus particle formation before release of the reproduced virus particles into the surrounding areas, such as the blood stream (Fielding et al., 2006; Kwon et al., 2020). Recent studies have revealed that the affinity of SARS-CoV-2 for ACE2 is 10-20-times higher than the affinity of SARS-CoV, which would explain its much higher transmissibility (Hoffmann et al., 2020; Zhang et al., 2020b). Upon binding, the viral S protein is subjected to proteolytic cleavage by the host cell’s transmembrane serine protease subtype 2 (TMPRSS2) (Hoffmann et al., 2020). The virus’ entry can be blocked by a clinically proven protease inhibitor, rendering the TMPRSS2 an interesting target against COVID-19 (Hoffmann et al., 2020). Interestingly, it was shown that SARS-CoV-2 does not use other receptors like aminopeptidase N or dipeptidyl peptidase 4 for cell entry as described for other coronaviruses (Zhou et al., 2020b). Therefore, these proteins are unlikely to represent suitable targets for therapy of COVID-19.

The tissue distribution of one of the main actors, ACE2, in organs like heart, kidney, endothelium, and intestine might explain the multi-organ dysfunction observed in COVID-19 patients (Zhang et al., 2020b). Several studies have provided information about the main symptoms, risk factors for severe disease progression, and clinical diagnostic values including blood routine, blood biochemistry, and infection-related biomarkers (Chen et al., 2020; Guan et al., 2020; Han et al., 2020; Zhou et al., 2020a). Although the details of these individual studies vary, there is a consensus among changes of numerous clinical parameters, which might be directly connected or must be considered together in a specific context. For example, hemoglobin is decreased in more than 50% of the patients, as is serum albumin in 98% (Chen et al., 2020; Guan et al., 2020; Yang et al., 2020; Zhou et al., 2020a). In intensive care unit (ICU) patients, reduced levels of hemoglobin levels and cluster of differentiation proteins (CD) 45, CD4, CD8, CD19, and CD16/56 were observed (Fan et al., 2020). In contrast, values of absolute lymphocyte count and absolute monocyte count were comparable to non-ICU patients (Fan et al., 2020). However, in contrast to human immunodeficiency virus (HIV) and cytomegalovirus, the CD4/8 ratio was not inverted (Fan et al., 2020). Main symptoms are fever, cough, and fatigue, all presenting reactions of an activated immune system (Guan et al., 2020). The activation of the immune and the complement system is also observed by a variety of markers including increased values for interleukin (IL)-6 (52% of the patients), erythrocyte sedimentation rate (85%), serum ferritin (63%), and C-reactive protein (CRP) (86%) (Chen et al., 2020; Risitano et al., 2020; Wang et al., 2020; Zhang et al., 2020c; Zhou et al., 2020a). Furthermore, recent studies that monitored and compared coagulation parameters of COVID-19 patients have suggested a tendency to procoagulant states (Han et al., 2020; Ji et al., 2020; Tang et al., 2020) and an increased risk of venous thromboembolism (Giannis et al., 2020), which was indicated by higher levels of fibrin/fibrinogen degradation products and fibrinogen itself, as well as lower antithrombin levels (Han et al., 2020). Moreover, D-dimer levels, a marker for coagulation and sepsis, are markedly increased in non-survivors of COVID-19 (Guan et al., 2020; Zhou et al., 2020a). Therefore, COVID-19 patients often suffer from leukopenia, lymphocytopenia, and thrombocytopenia (Guan et al., 2020).

Overall, these clinical parameters are interrelated when viewed from the perspective of heme and its interaction radius (Dutra and Bozza, 2014; Kühl and Imhof, 2014; Roumenina et al., 2016; Humayun et al., 2020). Heme is well-known as the prosthetic group of diverse proteins, e.g., hemoglobin, where it is responsible for the oxygen transport in the blood (Ascenzi et al., 2005). Under hemolytic conditions such as malaria, sickle cell anemia (SCD), β-thalassemia, and hemorrhage, or in case of severe cellular damage, heme is released in enormous amounts as a result of hemoglobin degradation, leading to a pool of labile heme (Ascenzi et al., 2005; Roumenina et al., 2016). In this case, the heme-detoxifying scavenger proteins, in particular hemopexin, become saturated, which allows heme to execute its wide-ranging effects (Chiabrando et al., 2014; Roumenina et al., 2016; Detzel et al., 2020). The response to heme in this context leads to cytotoxic, procoagulant, vasculotoxic, and proinflammatory effects, as well as an activation of the complement system (Dutra and Bozza, 2014; Roumenina et al., 2016). Labile heme also plays a central role in the pathology of severe sepsis which leads to vascular inflammation and severe toxic effects in organs like liver, kidney or cardiac tissue (Dutra and Bozza, 2014). These responses are, in part, mediated through direct interaction of heme with the responsible proteins (i.e. tumor necrosis factor α (TNFα), Toll like receptor 4 (TLR4), and complement factor 3 (C3) (Dutra and Bozza, 2014; Roumenina et al., 2016; Kupke et al., 2020), or by up-regulation of the respective cytokines, including IL-1β, IL-6, and TNFα (Dutra and Bozza, 2014). In addition, for heme a downstream ROS-dependent induction of distinct signaling pathways (MAPK/ERK pathway, NFκB signaling) is discussed, which can lead to stimulation of neutrophil recruitment, necrotic cell death or expression of adhesion molecules (Dutra and Bozza, 2014). In fact, we recently contextualized the role of heme as an inflammatory mediator as well as its crosstalk with the TLR4 signaling pathway (Humayun et al., 2020).

Although the aforementioned findings suggest a connection between biological processes implicated in SARS-CoV-2 and those related to heme, the lack of information on both subject areas impedes the use of modeling approaches that can facilitate their interpretation. Despite a considerable volume of research on SARS-CoV-2 over the past few months, knowledge of the molecular mechanisms responsible for the pathophysiology of the virus still remains scarce. Likewise, data in the context of heme’s biology is underrepresented, if available at all, in standard bioinformatics resources such as pathway databases. However, two of our recent scientific publications have focused on modelling knowledge around heme as well as SARS-CoV-2 (Domingo-Fernández et al., 2020; Humayun et al., 2020). Although both studies are tangential, each follows a knowledge-driven approach aimed at generating a context-specific Knowledge Graph (KG) that can subsequently be employed to investigate candidate mechanisms involved in hemolytic disorders and COVID-19. Moreover, in leveraging the interoperability between the COVID-19 and Heme KGs, these KGs can be integrated to shed light on the shared mechanisms between these two, seemingly independent domains.

Following the deposition of a manuscript stating that SARS-CoV-2 attacks hemoglobin 1-β chain and captures the porphyrin of its heme (Liu and Li, 2020), a heated debate arose about the scientific substantiation of the truthfulness of the claims. In this context, one should ask: How realistic is it, in view of the patients’ constitution that coronavirus components encounter heme and following up on this, what might be the consequences of such an interaction, if any? Thus, in order to contribute to the understanding of SARS-CoV-2, its strategies of infection, as well as the pathogenesis and the course of disease in coronavirus patients, we combined our know-how and expertise for a joint analysis of the aforementioned issue. On the one hand, we examine the possibility of a direct interaction of heme with select SARS-CoV-2 proteins and specific host cell proteins by applying our webserver HeMoQuest (Paul George et al., 2020) that is based on experimental data. One of the most promising findings was the prediction of heme-binding motifs (HBMs) in the host cell proteins ACE2 and TMPRSS2. On the other hand, by superimposing the two knowledge graphs, i.e. heme KG (Humayun et al., 2020) and COVID-19 KG (Domingo-Fernández et al., 2020), we provide insights into pathways that might play a role when considering heme in the context of COVID-19 infections. Finally, our results suggest that proinflammatory pathways could connect the pathophysiology of elevated heme with COVID-19 disease progression.

## 2 Materials and Methods

### 2.1 Screening for potential heme-binding motifs in COVID-19 related proteins

HeMoQuest (http://bit.ly/hemoquest) (Paul George et al., 2020) was used to identify potential HBMs in the following proteins of SARS-Cov-2: S protein, M protein, E protein, and protein 7a, and human: ACE2 and TMPRSS2. For the selection, the procedure described earlier was applied (Wißbrock et al., 2019; Detzel et al., 2020).

### 2.2 Homology modeling

In all *in silico* experiments, we used available cryogenic electron microscopy (cryoEM) structures or homology models (publicly available or in-house built). In case of ACE2 the recently published, fully glycosylated cryoEM structure (PDB: 6M18) was used, which was recorded in complex with sodium-dependent neutral amino acid transporter B (B^0^AT1) (Yan et al., 2020). B^0^AT1 was removed in order to focus on ACE2 only. Although S protein cryoEM structures for open and closed states are available (PDB: 6VXX and 6VYB) (Walls et al., 2020), these structures lack several surface-exposed sequence stretches, in which some of the predicted motifs are located. We therefore used the structure available from the C-I-TASSER structure prediction server reported earlier (Zhang et al., 2020a). For SARS-CoV-2 protein 7a and TMPRSS2, no structures are available so far. Thus, homology models were built using YASARA version 19.12.14 and the *hm_build* macro with default settings (Krieger and Vriend, 2014). We were able to build a hybrid model of the virion surface-exposed part of protein 7a from two structures of the SARS virus protein 7a (PDB: 1XAK and 1YO4). The model achieved an overall Z-score of −0.053, which can be regarded as suitable. In contrast, a hybrid model of TMPRSS2 based on kallikrein and hepsin (PDB: 6I44, 6O1G, 1Z8G, 5CE1) exhibited only a poor overall Z-score of −2.363 and was therefore rejected. In this case, further *in silico* analysis was performed with the available Swiss-model O15393 (Waterhouse et al., 2018).

### 2.3 Modeling the interplay between SARS-CoV-2 and heme

In order to investigate the mechanisms linking SARS-CoV-2 and heme, we exploited the KGs generated in our previous work (Domingo-Fernández et al., 2020; Humayun et al., 2020). We compiled the two KGs encoded in Biological Expression Language (BEL) using PyBEL (Hoyt et al., 2018) directly from their public repositories (i.e. https://github.com/covid19kg and https://github.com/hemekg/) and superimposed their interactions onto a merged network. Given the high degree of expressivity of BEL that enables the representation of multimodal biological information, the KGs were not only enriched with molecular information, but also with interactions from the molecular level to phenotypes and clinical readouts. We leveraged this multimodal information to hypothesize the pathways that connect key molecules associated with SARS-CoV-2 and heme to phenotypes observed in COVID-19 patients.

Since both KGs comprise several thousands of interactions, manually inspecting all relations and evaluating the implication of the crosstalk between COVID-19 and heme is largely infeasible. Accordingly, this analysis primarily focuses on the set of nodes present in both KGs. Prior to the crosstalk analysis, we conducted a one-sided Fisher’s exact test (Fisher, 1992) to evaluate the significance of the overlap between human proteins present in each of the KGs (*p*-value < 0.01). We then classified the set of overlapping nodes into four pathways based on their functional role: i) immune response – inflammation, ii) immune response – complement system, iii) blood and coagulation system, and iv) organ-specific diagnostic markers. Finally, for each of the pathways, we analyzed the similarity between the signatures for both KGs by superimposing the relations that connect each of the overlapping nodes to COVID-19 and heme biology. The relations present in Figure 2 are also shown in Table S1-S4 together with their evidence and provenance information.

**Fig. 1:**
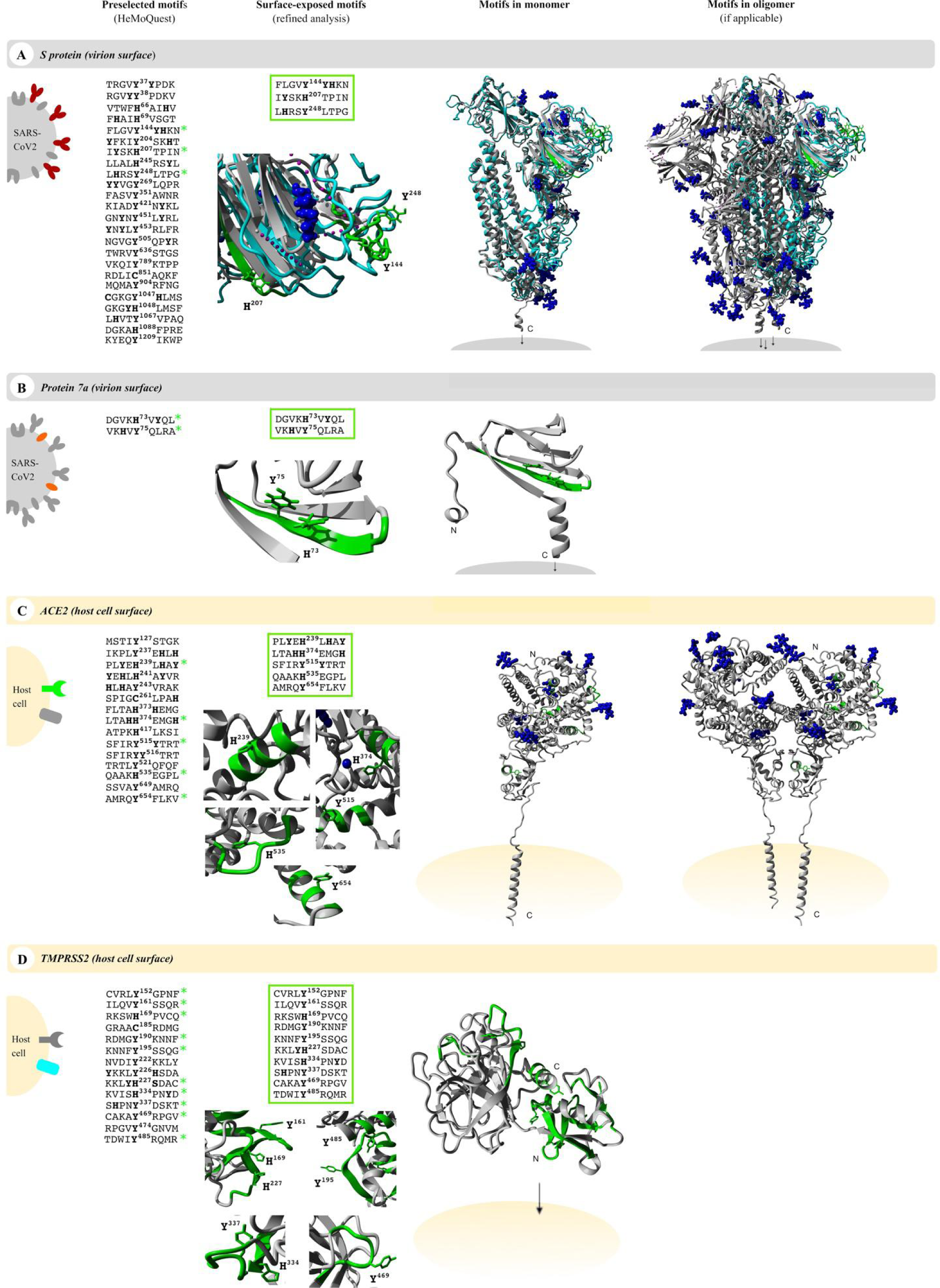
Potential heme-binding proteins on the virus and host cell surface. Four COVID-19-related proteins, namely the virus proteins (grey) **(A)** S protein and **(B)** protein 7a as well as the host cell proteins (yellow) **(C)** ACE2 and **(D)** TMPRSS2, turned out as possible heme-binding proteins. The location of the proteins is presented (first column, left), individually highlighting each target protein (S protein, red; protein 7a, orange; ACE, green; TMPRSS2, turquoise). All motifs predicted by HeMoQuest (Paul George et al., 2020) are shown excluding those with modifications (glycosylation, disulfide bonds) or located in intracellular or virion domains, i.e. 24 motifs for S protein, 2 motifs for protein 7a, 15 motifs for ACE2, and 14 motifs for TMPRSS2. Potential heme-binding residues are bold-written and numbered according to SwissProt numbering system (Bairoch and Apweiler, 2002). A refined analysis considering the surface accessibility of the motifs resulted in 3 motifs for S protein, 2 largely overlapping motifs for protein 7a, 5 motifs for ACE2, and 10 motifs for TMPRSS2 (third column). In addition, these motifs are highlighted in a zoom-in below the list with annotation of the respective potential coordinating residues (green; third column), as well as in the available monomer (fourth column) and oligomer (fifth column) structures, if applicable (S protein, homology model from C-I-TASSER (Zhang et al., 2020a); protein 7a, in-house homology model; ACE2, PDB: 6M18; TMPRSS2, Swiss-model: O15393). Within the oligomers, the motifs were only depicted in one of the monomers (green). Each time, the central, potential heme-coordinating residue is shown as stick model. Since some surface-exposed motifs within S protein were not covered by the available EM structure (PDB: 6VXX), motifs were highlighted within the monomer homology model from C-I-TASSER (Zhang et al., 2020a) (turquoise), which was then superimposed with the trimer (PDB: 6VXX). Where applicable, glycosylation sites and ions are highlighted in blue.

**Fig. 2:**
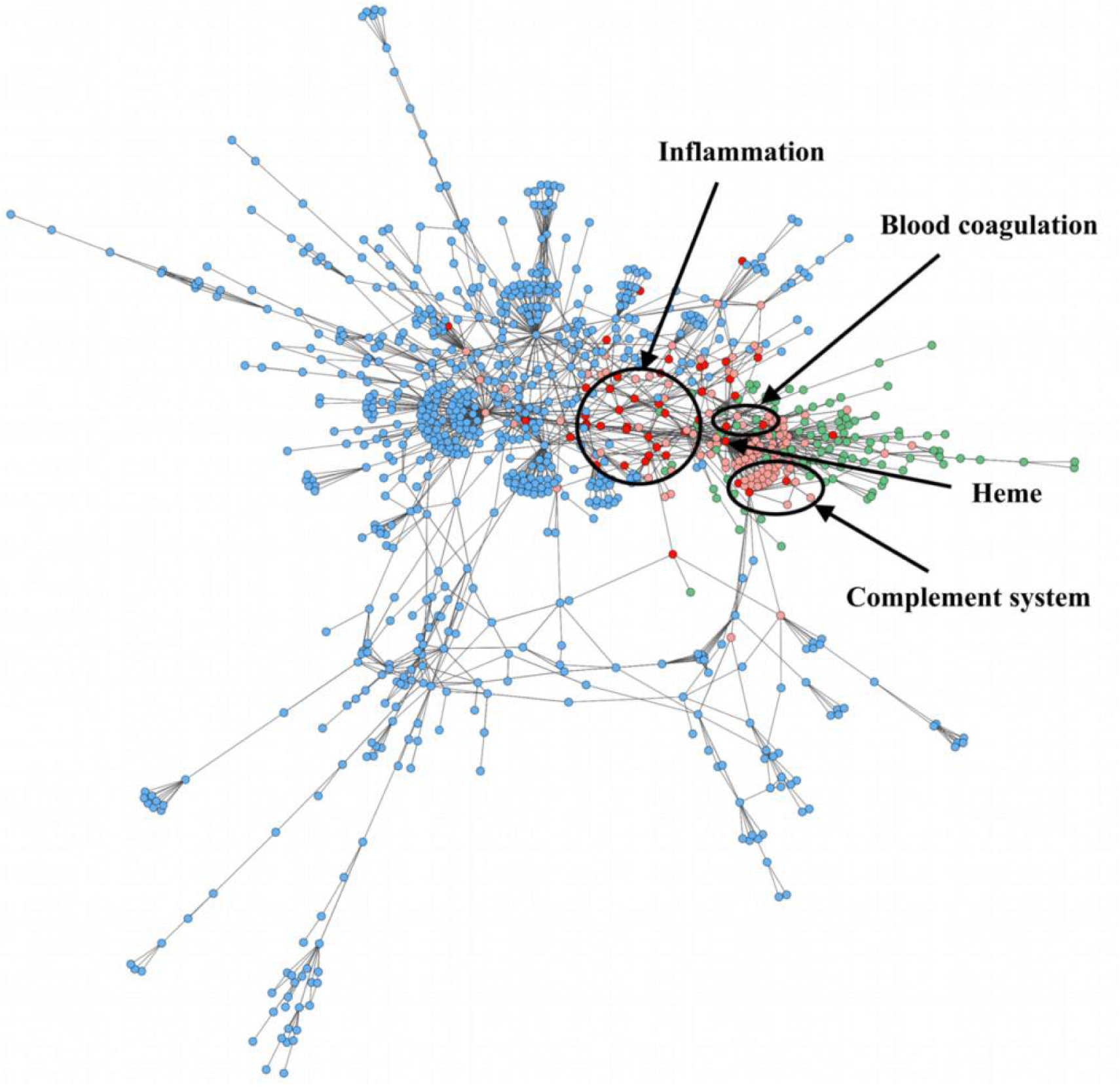
Overlap between the COVID-19 KG (Domingo-Fernández et al., 2020) and Heme KG (Humayun et al., 2020). The figure depicts the largest component of the overlap between the two KGs exclusively comprising molecular interactions (i.e., only relations comprising proteins and small molecules are shown). The color of the nodes denotes whether it is present exclusively in the COVID-19 KG (blue), in Heme KG (green), or in both (red). Finally, to highlight the areas close to the overlapping nodes, their neighbors are colored in light red. The most matching nodes are shown circled assigned to the respective major systems, i.e. inflammation, blood coagulation and complement system.

In order to validate the knowledge-driven hypothesis coming from the KG, we compared the relations emerging from the overlap between the two KGs with experimental data published in the context of COVID-19 (Blanco-Melo et al., 2020). The concordance of the expression patterns in these datasets with each relation shown in Figure 3 is shown in Table S5.

**Fig. 3:**
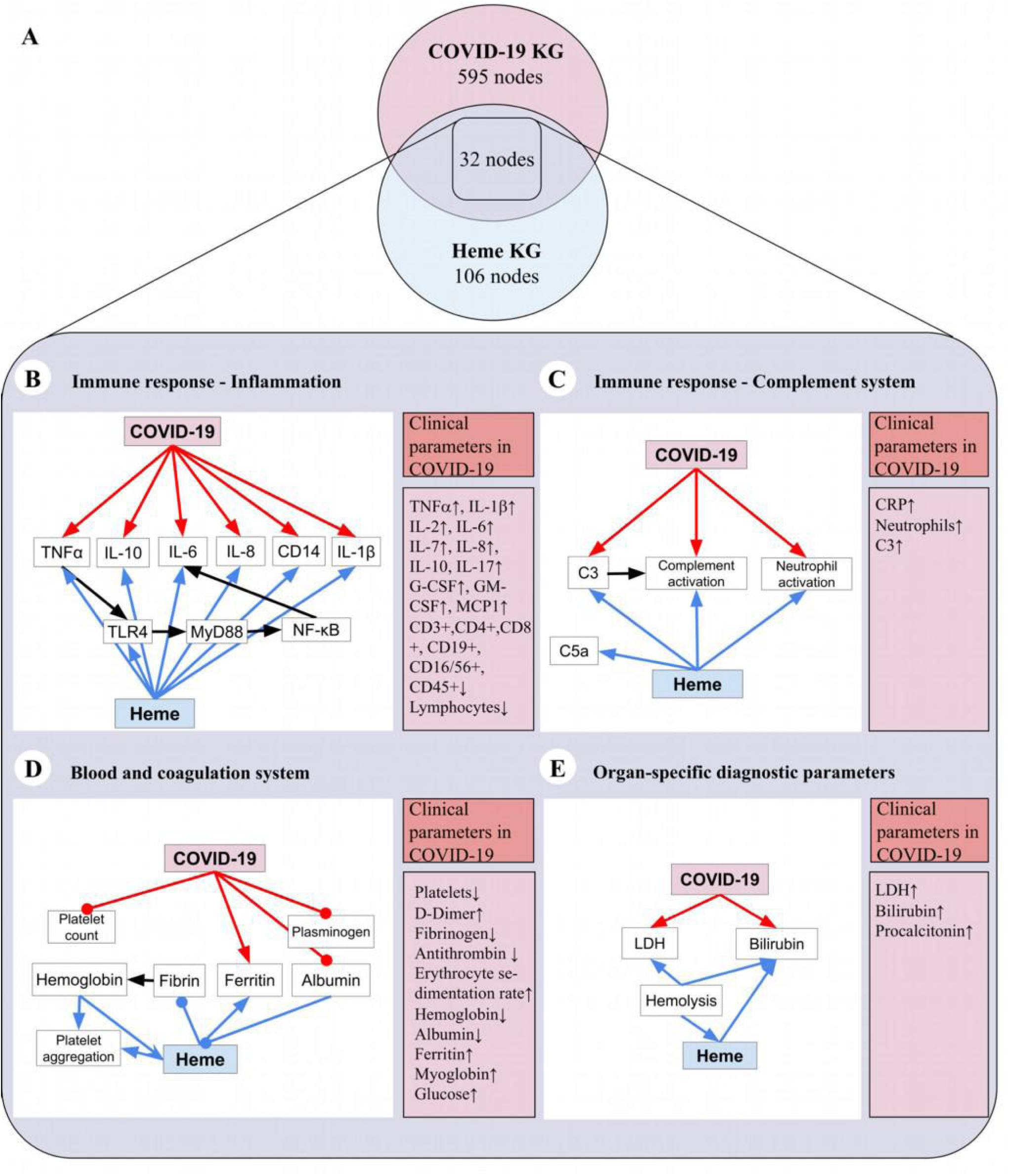
Shared biochemical pathways based on the overlap between COVID-19 KG (Domingo-Fernández et al., 2020) and Heme KG (Humayun et al., 2020). **(A)** Overlap between the two KGs based on human proteins represented as a Venn Diagram. The numbers of nodes that only present proteins are depicted (595 nodes in COVID-19 KG, 106 nodes in Heme KG and 32 nodes as an overlap between both KGs). In particular, the two KGs overlap in the following systems: **(B)** Immune response – inflammation, **(C)** immune response – complement system, **(D)** blood and coagulation system, and **(E)** organ-specific diagnostic markers. Moreover, for each classification available clinical parameters were denoted (Chen et al., 2020; Risitano et al., 2020). CRP = C-reactive protein, C3 = Complement component 3, CD = Cluster of differentiation, G-CSF = Granulocyte-colony stimulating factor, GM-CSF = Granulocyte macrophage colony-stimulating factor, IL = Interleukin, LDH = Lactate dehydrogenase, MCP1 = Monocyte chemoattractant protein 1.

## 3 Results

COVID-19 progression severely diverges between affected patients with ARDS and other patients, which could even remain asymptomatic. Current research is thus focusing on explaining the reasons for such discrepancy considering the physical conditions and (pre-)existing illnesses of those affected. With regard to the subject of a possible interrelation between COVID-19 and heme, numerous options need to be regarded. First, the earlier claim of an interaction of protoporphyrin IX with SARS-CoV-2 (Liu and Li, 2020) must be questioned, since heme would appear before protoporphyrin IX as a consequence of e.g., hemolytic conditions. Thus, the direct interaction of heme with viral surface proteins, as well as host cell proteins exposed to virus attack, needs to be considered. Second, systemic hyperinflammation follows severe COVID-19 infection. This is manifested by an increase in the abundance of numerous cytokines (e.g., IL-2, IL-7, IL-6, TNFα) (Ye et al., 2020), which is indicative of cytokine release syndrome (Huang et al., 2020) and leads to elevated serum biomarkers in patients (e.g., CRP, lactate dehydrogenase (LDH), D-dimer, ferritin) (Chen et al., 2020; Young et al., 2020; Zhang et al., 2020c; Zhou et al., 2020a). Several of these indications, however, were also reported for labile heme occurring in patients with hemolytic disorders (Litalien et al., 1999; Barcellini and Fattizzo, 2015). Therefore, heme as a key player in initiating or mediating distinct processes in connection with a viral infection needs to be considered as well. This can be exemplified with the interaction of heme with e.g., Zika, Chikungunya, and HIV-1 viruses (Gupta et al., 2015; Lecerf et al., 2015; Assunção-Miranda et al., 2016; Neris et al., 2018). In the following, we present our results concerning the potential direct heme interaction with COVID-19-related proteins as well as a detailed analysis of common pathways of excess heme and COVID-19 pathophysiology.

### 3.1 Heme-binding ability of surface proteins of SARS-CoV-2 and host cells

Numerous interesting target proteins of the virus and host cell surface were linked with pathological effects of SARS-CoV-2, including E protein, S protein, M protein and protein 7a as well as the human proteins ACE2 and TMPRSS2. All proteins contain at least an extracellular, surface-exposed part and are thus accessible for interaction with heme (Hänel and Willbold, 2007; Mendes de Oliveira et al., 2018; Mousavizadeh and Ghasemi, 2020; Walls et al., 2020). This led us to examine these proteins for potential heme-binding sites. We identified potential HBMs on all target proteins using the machine-learning algorithm HeMoQuest (Paul George et al., 2020). Screening of the amino acid sequences of S protein, protein 7a, ACE2 and TMPRSS2 resulted in 50, 6, 21, and 32 potential HBMs, respectively. M protein and E protein were dismissed as candidates, since no suitable HBMs were found. HBMs, which are part of the transmembrane or intravirion/intracellular domains, were removed from the selection. In addition, we excluded motifs in which the potential coordinating residue or a residue adjacent to the coordination site was involved in disulfide bonds or glycosylation. After this refinement of the hits, we identified 24 motifs in S protein, two in protein 7a, 15 in ACE2, and 14 in TMPRSS2 (Figure 1). These motifs were then manually screened for surface accessibility using the protein structures, or if unavailable, homology models. Consequently, three motifs for S protein, two motifs for protein 7a, five motifs for ACE2 and ten motifs for TMPRSS2 remained and are discussed below (Figure 1). The potential HBMs in S protein are all located in the N-terminal domain (Figure 1A) (Ou et al., 2020). The first occurring sequence FLGV**Y**^144^**YH**KN may be the most promising HBM, which is based on a YYH motif and further equipped with phenylalanine at P-4, two additional hydrophobic amino acids (Val, Leu), and a net charge of +2, all beneficial for heme binding (Syllwasschy et al., 2020). The following, I**Y**SK**H**^207^TPIN and L**H**RS**Y**^248^LTPG, contain a Y/H-based motif with two spacers between the potential coordinating residues, e.g., YXXH, which have been shown to be less favorable for heme binding (Syllwasschy et al., 2020). Nonetheless, both motifs possess a net charge of +2, and several hydrophobic residues, and are thus likely to moderately bind heme. In protein 7a, only two overlapping motifs were predicted, which is not surprising due to the small size of 121 amino acids (Figure 1B). Both, DGVK**H**^73^V**Y**QL and VK**H**V**Y**^75^QLRA, possess a HXY motif (Syllwasschy et al., 2020) and three hydrophobic residues, rendering it a moderate heme binder and, in turn, protein 7a as a less interesting candidate for interaction with heme.

The analysis of ACE2 revealed five HMBs in total, two of which representing promising H/Y motifs (Figure 1C). The most interesting HBM is LTA**HH**^374^EMG**H** comprising a HXXXH motif, which was recently shown to exhibit high heme-binding affinity (Syllwasschy et al., 2020). The central H^374^ is immediately adjacent to the site that is essential for cleavage by ADAM17 (Heurich et al., 2014; Lan et al., 2020). The occurrence of three histidines may be favorable for heme binding as could L^370^, while E^375^ might be slightly detrimental. The second interesting motif is PL**Y**E**H**^239^L**H**A**Y**, since it contains the efficient HXH motif (Syllwasschy et al., 2020) with further advantageous aromatic tyrosines (Y^237^, Y^243^) and hydrophobic leucines (L^236^, L^240^). The only limitation to affinity might be E^238^. The remaining three motifs SFIR**Y**^515^**Y**TRT, QAAK**H**^535^EGPL, and AMRQ**Y**^654^FLKV are less promising because they only contain one coordinating amino acid or the weak motif YY^80^.

The largest number of motifs, i.e. ten in total, was identified in the transmembrane serine protease TMPRSS2. Four of these motifs contained only one coordinating amino acid and can be dismissed for the aforementioned reasons. Additional three motifs (CVRL**Y**^152^GPNF, RKSW**H**^169^PVCQ, CAKA**Y**^469^RPGV) contain cysteine as a possible further coordinating site. Cysteine has been shown to efficiently function as HRM in conjunction with proline in the CP motif, but without it, it lacks high heme-binding affinity (Brewitz et al., 2015). Two further overlapping motifs (KVIS**H**^334^PN**Y**D and S**H**PN**Y**^337^DSKT) were found in the protease domain of TMPRSS2 (Figure 1D). Both are equally well-suited for moderate heme binding based on the HXXY motif and a positive net charge. Within the scavenger receptor cysteine-rich domain of the enzyme (Mendes de Oliveira et al., 2018), the interesting motif KKL**YH**^227^SDAC was found. It features an YH motif of intermediate heme-binding affinity on peptide level, however, of markedly improved affinity on the protein level as earlier demonstrated for IL-36α (Wißbrock et al., 2019; Syllwasschy et al., 2020). Furthermore, it shows high net charge and a hydrophobic leucine, which likely leads to high heme-binding affinity. A comparative analysis of all motifs revealed that the S protein was the only SARS-CoV-2-derived protein with a promising HBM (FLGV**Y**^144^**YH**KN). In contrast, the human proteins ACE2 and TMPRSS2 showed both quantitatively and qualitatively superior motifs. In ACE2, LTA**HH**^374^EMG**H** and PL**Y**E**H**^239^L**H**A**Y** boast promising H/Y motifs and highly favorable properties. Likewise, TMPRSS2 contains the motifs KVIS**H**^334^PN**Y**D and S**H**PN**Y**^337^DSKT, which represent potential heme-interaction sites directly in the catalytic protease domain, and KKL**YH**^227^SDAC, which might also bind heme efficiently.

### 3.2 Pathophysiological effects of heme and COVID-19 intersect at inflammation

In order to shed light on the crosstalk and common pathways between heme and COVID-19, we investigated the overlap between our two KGs (i.e., heme KG (Humayun et al., 2020) and the COVID-19 KG (Domingo-Fernández et al., 2020)) (Figure 2). While the Heme KG was generated from the analysis of 46 scientific articles specifically selected to explain inflammatory processes related to labile heme, the COVID-19 KG contains over 150 articles. The difference in the size of these KGs thus explains the disproportionate number of molecules they possess. Nonetheless, we observed that a significant amount of proteins is shared, predominantly in three major systems, namely blood coagulation, complement and immune system. Among the 85 shared nodes, there are 45 clinical phenotypes, 35 proteins, 4 immune system specific cells, and 5 small molecules. 27 nodes belong to immune response evoking (pro-)inflammatory pathways, 4 to the complement system, and 22 to the blood coagulation system (Figure 3A). Moreover, we also noticed the presence of 7 clinical phenotypes related to organ dysfunction. Further, we individually investigated the four systems to reveal the common relations observed in each of the two KGs (Figure 3, Table S1-S4). Finally, we evaluated the concordance of these systems with experimental data published in the context of COVID-19 (Blanco-Melo et al., 2020) and the vast majority of them are in line with the findings presented below (Table S5).

The largest consistency was found in inflammatory pathways (Figure 3B) as indicated by a common set of inflammatory – mostly pro-inflammatory – molecules. These are changed with respect to their levels due to expression and/or secretion or their activity as a consequence of both, high heme concentrations and COVID-19 infection, mediating inflammatory response. In particular, the pro-inflammatory cytokines TNFα, IL-1β, IL-6, IL-8, and the anti-inflammatory cytokine IL-10, as well as proteins related to TLR4-mediated signaling pathways (i.e. CD14, MyD88, NF-κB and TLR4) are influenced under both conditions (Figure 3B, Table S1).

Within the complement system (Figure 3C, Table S2), one of the main mediators, C3, is activated under hemolytic conditions associated with high heme concentrations, thus leading to complement activation (Roumenina et al., 2016). The same was observed in COVID-19 patients (Risitano et al., 2020). Furthermore, other complement factors, like C5a and C1q, were reported to be activated by heme (Roumenina et al., 2016). So far, an increase of the activation of these proteins was not described for COVID-19. Finally, the number of neutrophils is positively correlated with both heme and COVID-19 infection. However, heme induces neutrophil activation through a ROS-dependent mechanism (Dutra and Bozza, 2014), a pathway that is not yet discussed in the context of COVID-19.

The blood and coagulation system is pronounced by the connecting proteins ferritin and albumin (Figure 3D, Table S3). Both conditions lead to reduced levels of ferritin, a protein involved in iron uptake and release (Mogl, 2007). Same applies for albumin in COVID-19 patients (Chen et al., 2020; Guan et al., 2020; Yang et al., 2020; Zhou et al., 2020a). Moreover, albumin is known as one of the common heme scavengers, neutralizing heme’s toxic effects up to a certain extent (Roumenina et al., 2016). As indicated by the impact on different components of the blood coagulation system, such as plasminogen or fibrin in case of COVID-19 and heme, respectively, both conditions can influence hemostasis. With regard to the impact on platelets, a decreased platelet count was observed in COVID-19 patients (Wang et al., 2020), whereas for heme an induction of platelet aggregation was described (Roumenina et al., 2016).

Finally, a trend towards elevated levels of organ-specific diagnostic markers, i.e. LDH and bilirubin, is shared by both KGs (Figure 3E, Table S4).

## 4 Discussion

Currently, SARS-CoV-2 and its associated disease COVID-19 keep the world in suspense. Patients being most severely affected suffer from pneumonia, acute respiratory distress syndrome, and death (Ye et al., 2020; Zhou et al., 2020a). While COVID-19 patients often exhibit high levels of proinflammatory markers as well as an activation of the complement and coagulation system, hemoglobin and albumin levels have been reported to be remarkably low (Chen et al., 2020; Risitano et al., 2020; Ye et al., 2020). These affected clinical parameters have generated a recent debate about the role of heme in the context of COVID-19 that has not been conclusively explained to date (Liu and Li, 2020).

With this work, we intend to provide deeper insights into the connection between SARS-CoV-2 infection, COVID-19 and the effects of heme, wherever possible and appropriate. Such a connection would be in line with recent studies that already described the impact of heme in the context of different viruses (Lecerf et al., 2015; Gupta et al., 2015; Neris et al., 2018). Lecerf et al. reported on the interaction of heme with antibodies (Abs) resulting in the induction of new antigen binding specificity and acquisition of binding polyreactivity to gp120 HIV-1 in 24% of the antibodies from different B-cell subpopulations of seronegative individuals (Lecerf et al., 2015). In contrast, no difference in the sensitivity towards heme was found for Abs originally expressed by naive, memory, or plasma cells. The transient interaction of heme with a fraction of circulating Abs that might change their antigen binding repertoire by means of cofactor association was suggested as another possible regulatory function of heme (Lecerf et al., 2015). In addition, the novel antigen specificities of these circulating Abs was proposed to be recruited only in case of certain pathological conditions that might depend on extracellular heme as occurring in disorders such as malaria, sickle cell disease, hemolytic anemia, β-thalassemia, sepsis, and ischemia-reperfusion (Lecerf et al., 2015). A similar report by Gupta et al. revealed the heme-mediated induction of monoclonal immunoglobulin G1 antibodies that acquired high-affinity reactivity towards antigen domain III of the Japanese encephalitis virus (JEV) E glycoprotein that exhibited neutralizing activity against dominant JEV genotypes (Gupta et al., 2015). In both cases, heme was found to confer novel binding specificities to the respective Abs without changing the binding to their cognate antigen and, as a consequence of the contact with heme, the anti-inflammatory potential of these Abs was substantially increased (Gupta et al., 2015). Finally, Assuncao-Miranda et al. and Neris et al. described the inactivation of different arthropod-borne viruses like Dengue, Yellow Fever, Zika, Chikungunya, Mayaro and others by porphyrin treatment via targeting of the viral envelope and thus, the early steps of viral infection (Assunção-Miranda et al., 2016, Neris et al., 2018). All together, these studies advocate for studying the impact of heme in coronavirus-infected patients (Figure 4).

**Fig. 4:**
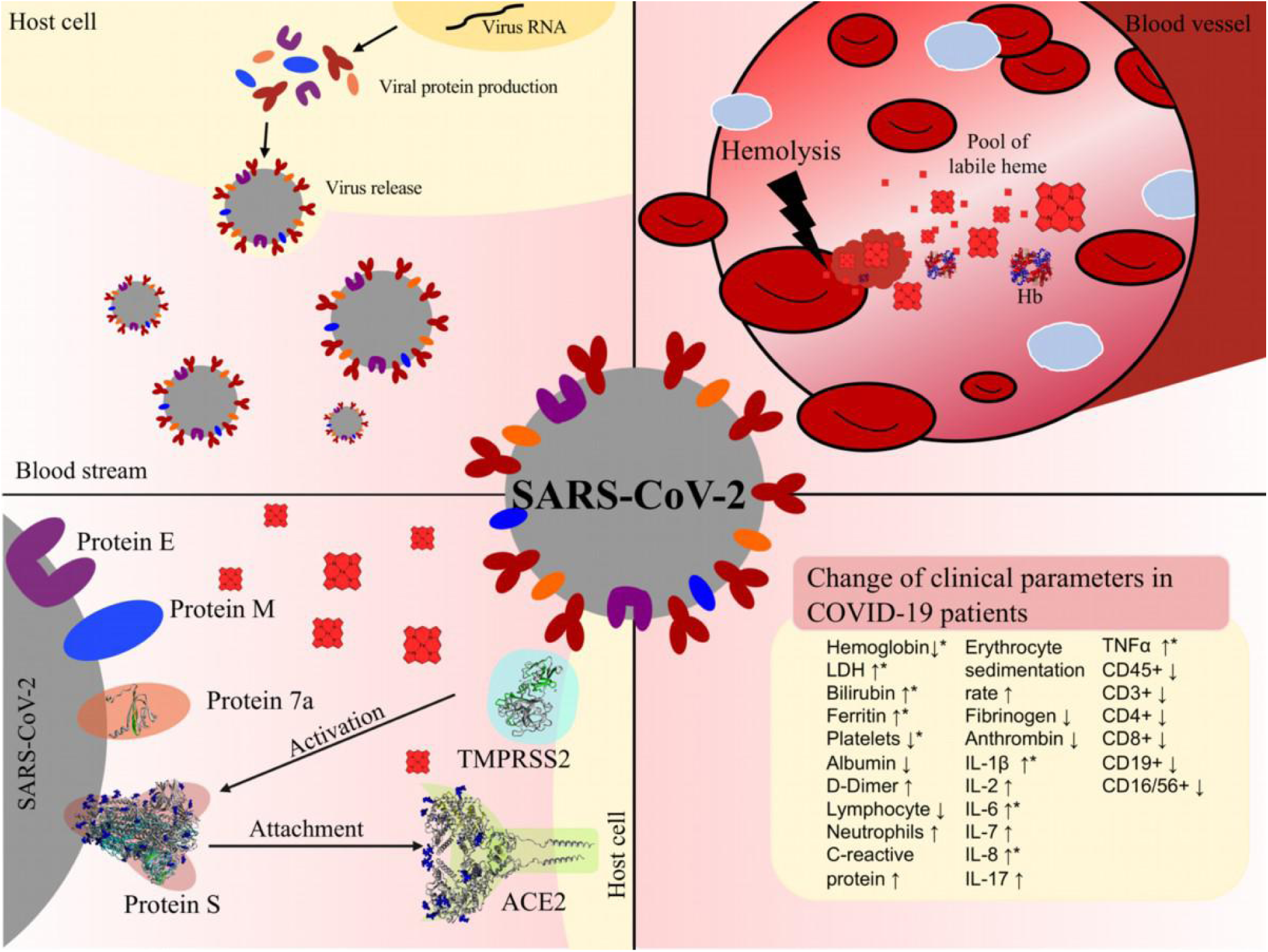
COVID-19 infection and hemolysis show common changes of clinical parameters. Top left: Virus (grey) release after conquering the host cell (yellow) and taking over its protein synthesis machinery. Top right: In case of hemolysis, erythrocyte lysis occurs in a blood vessel, leading to degradation of hemoglobin (blue/red, PDB: 1GZX) and, thus, to an excess of labile heme. Circulating erythrocytes (red) and platelets (bright blue) are shown. Bottom left: Interaction of virus and host cell before cell entry. S protein (light red, PDB: 6VXX) on the surface of SARS-CoV-2 (grey), interacts with human ACE2 receptor (green, PDB: 6M18). The protease TMPRSS2 (turquoise) primes S protein and contributes to cell entry (Hoffmann et al., 2020). The accessory protein 7a (in-house homology model) interacts with S protein, M protein and E protein for virus particle assembly. E protein is discussed to form ion channels and to play a role in viral genome assembly (Ruch and Machamer, 2012). M protein may be relevant for entry and attachment of the virus as well as for the budding process (Bianchi et al., 2020). The shown proteins contain domains in the extracellular space that have specific characteristics for heme binding. Bottom right: Prominent changes of clinical parameters in patients suffering from COVID-19 infection (↑ increase, ↓ decrease). The terms depicted by an asterisk, have been reported in both, hemolysis and COVID-19 infection (Barcellini and Fattizzo, 2015; Chen et al., 2020).

Here, we have investigated the possibility of a direct interaction of heme with SARS-CoV-2 surface proteins and their human counterparts ACE2 and TMPRSS2. Our analysis revealed that heme binding conferred by HBMs would potentially be possible. The quality, availability, and accessibility of the motifs follows the rank order: TMPRSS2 (good binder) > ACE2 > S protein > Protein 7a (poor binder). Especially in TMPRSS2, the location of the most suitable HBMs correlates with the important catalytic protease domain. This potential heme interaction would be of a transient nature, as has been observed for other heme-binding proteins such as IL-36α and CBS (Kumar et al., 2018; Wißbrock et al., 2019). Intact heme would bind to the protein surface in a reversible fashion, which would be in contrast to the recently presented hypothesis by Liu & Li (Liu and Li, 2020). Therein, the authors describe heme extraction from hemoglobin through attack by viral proteins and subsequent iron and porphyrin release from heme, which does not occur in a physiological situation (Belcher et al., 2010). In addition, the docking analysis performed in their study is not based on experimental data concerning the porphyrin interaction, unlike the data-driven machine learning algorithm HeMoQuest (Paul George et al., 2020) used in our study. Nevertheless, the effect of heme on the suggested proteins TMPRSS2, ACE2, S protein, and Protein 7a needs experimental verification.

Apart from investigating the direct impact of heme on proteins at the interface of the virus-host cell interaction, we also explored similarities between relevant pathways characterizing the respective pathologies, i.e. labile heme occurrence in hemolytic conditions and COVID-19 disease progression (Figure 4). Both, hemolytic conditions and COVID-19, have been found to trigger inflammatory pathways. COVID-19 patients often develop respiratory distress syndrome, which is accompanied by a cytokine storm, and thus an activation of the immune system (Ye et al., 2020). Clinically, this is manifested by an increase in the levels of a wide range of cytokines, including TNFα, IL-1β, IL-6 and IL-8 (Ye et al., 2020), and the activation of the complement system (e.g. C3) (Risitano et al., 2020). Interestingly, hemoglobin is described to be often decreased in COVID-19 patients without indicating the molecular cause (Huang et al., 2020). This seems to correlate with increased levels of the iron-storage protein ferritin. An increase in ferritin concentration is observed in diseases like hemochromatosis or porphyria (Mogl, 2007). Furthermore, it is upregulated during hemolytic diseases as a consequence of hemoglobin degradation and the associated increase of oxidative stress, e.g. induced by heme (Belcher et al., 2010). Hemolytic disorders such as malaria, ischemia-reperfusion, hemorrhage or hemolytic anemias are associated with an excess of labile heme and are, as in COVID-19 infection, often accompanied by inflammatory events (Chiabrando et al., 2014; Barcellini and Fattizzo, 2015). Therefore, similar clinical parameters are observed under these conditions (Barcellini and Fattizzo, 2015). Moreover, several studies have reported that heme directly binds or induces TNFα, IL-1β and IL-8, and triggers numerous inflammatory pathways (e.g. NF-κB signaling) (Dutra and Bozza, 2014; Humayun et al., 2020). Taken together, these clinical observations suggested a correlation between both processes, which we aimed to analyze by superimposing the two KGs of both pathophysiologies (Domingo-Fernández et al., 2020; Humayun et al., 2020). Indeed, the results of the knowledge-driven analysis revealed a core of similar molecular patterns shared. The majority of these were related to the three major systems inflammation, complement, and coagulation system. As expected, inflammation was the most emphasized common system, suggesting several processes that are commonly mediated by heme and in COVID-19. The TLR4 signaling pathway was previously shown to play an important role in heme-mediated inflammatory processes. Interestingly, this pathway with its components TLR4, MyD88 and NF-κB was pronounced in the overlay of the heme KG and COVID-19 KG. The TLR4 pathway belongs to the innate immune system, and thus results in the production of several proinflammatory cytokines, such as TNFα, IL-1β and IL-6 (Humayun et al., 2020). TNFα and IL-1β can further stimulate the release of inflammatory mediators, such as IL-8. Exactly the same proteins have emerged as common key molecules in our analysis. Clinical observations revealed their upregulation in COVID-19 patients as well as during hemolytic events (Dutra and Bozza, 2014; Humayun et al., 2020; Ye et al., 2020), which highlights even more the TLR4 signaling pathway in both situations. Interestingly, TNFα and IL-1β were reported to be capable of regulating platelet aggregation. This might support the common link of both pathologies to blood coagulation (Bar et al., 1997). Blood parameters, such as hemoglobin and albumin levels, may allow for a direct correlation between COVID-19 and heme, since they are inevitably connected to the processing of heme under hemolytic conditions (Chiabrando et al., 2014; Roumenina et al., 2016). At the current state of research, there is no explanation for the decreased levels of hemoglobin in COVID-19. It might be conceivable, that this is due to a rapid turnover of red blood cells, which would lead to a degradation of hemoglobin and, in turn, to an increase of heme. Although for some viruses it is known that they cause hemolysis of red blood cells, such as hepatitis A (Goel et al., 2018), a similar behavior has not yet been described for SARS-CoV-2 and related viruses.

However, our approach is not without limitations as our analysis is restricted to a limited number of scientific articles. Furthermore, there is an unbalanced source of information when comparing the tremendous amount of literature that is currently being published on COVID-19 versus the information that is currently included in heme KG for heme. The findings described herein require a more detailed experimental investigation with dedicated experiments for each of the reported relations that shed light on the underlying biochemical mechanisms as well as for the full characterization of the heme-binding capacity of the proposed proteins. Nevertheless, the results of this study draw attention to a relationship that could be plausible based on the current characterization of COVID-19 by clinical parameters. A correlation between the symptoms of COVID-19 infection and the consequences of excess heme does not necessarily have to be related, but in specific cases it may correlate or even cause a more severe course of the disease in existing hemolytic conditions or hemolysis-provoking events.

## 6 Author Contributions

M.H.A. and D.I. designed and planned the project. M.-T.H., D.D.-F., Y.G., M.S.D, and B.F.S. performed the experiments and collected the data. Data analysis was carried out by all authors. The manuscript was written through the contribution of all authors.

## 7 Acknowledgments

This work has been supported by the MAVO and ICON programs of the Fraunhofer Society. Financial support by the University of Bonn is gratefully acknowledged.

## References

Alsaadi, E. A. J., and Jones, I. M. (2019). Membrane binding proteins of coronaviruses. Future Virol. 14, 275–286. doi: 10.2217/fvl-2018-0144.

Ascenzi, P., Bocedi, A., Visca, P., Altruda, F., Tolosano, E., Beringhelli, T., et al. (2005). Hemoglobin and heme scavenging. IUBMB Life 57, 749–759. doi: 10.1080/15216540500380871.

Assunção-Miranda, I., Cruz-Oliveira, C., Neris, R. L. S., Figueiredo, C. M., Pereira, L. P. S., Rodrigues, D., Da Poian, A. T., Bozza, M. T. (2016). Inactivation of Dengue and Yellow Fever viruses by heme, cobalt-protoporphyrin IX and tin-protoporphyrin IX. J. Appl. Microbiol. 120, 790–804. doi: 10.1111/jam.13038.

Bairoch, A., and Apweiler, R. (2002). The SWISS-PROT protein sequence database and its supplement TrEMBL in 2000. Nucleic Acids Res. 28, 45–48. doi: 10.1093/nar/28.1.45.

Bar, J., Zosmer, A., Hod, M., Elder, M. G., and Sullivan, M. H. (1997). The regulation of platelet aggregation in vitro by interleukin-1beta and tumor necrosis factor-alpha: changes in pregnancy and in pre-eclampsia. Thromb. Haemost. 78, 1255–1261.

Barcellini, W., and Fattizzo, B. (2015). Clinical applications of hemolytic markers in the differential diagnosis and management of hemolytic anemia. Dis. Markers 2015, 635670. doi: 10.1155/2015/635670.

Belcher, J. D., Beckman, J. D., Balla, G., Balla, J., and Vercellotti, G. (2010). Heme degradation and vascular injury. Antioxid. Redox Signal. 12, 233–248. doi: 10.1089/ars.2009.2822.

Bianchi, M., Benvenuto, D., Giovanetti, M., Angeletti, S., and Pascarella, S. (2020). Sars-CoV-2 Envelope and Membrane proteins: differences from closely related proteins linked to cross-species transmission? Preprints. doi: 10.20944/preprints202004.0089.v1.

Blanco-Melo, D., Nilsson-Payant, B. E., Liu, W.-C., Uhl, S., Hoagland, D., Møller, R., et al. (2020). Imbalanced host response to SARS-CoV-2 drives development of COVID-19. Cell 181, 1036–1045.e9. doi: 10.1016/j.cell.2020.04.026.

Brewitz, H. H., Kühl, T., Goradia, N., Galler, K., Popp, J., Neugebauer, U., et al. (2015). Role of the chemical environment beyond the coordination site: Structural insight into Fe(III)protoporphyrin binding to cysteine-based heme-regulatory protein motifs. ChemBioChem 16, 2216–2224. doi: 10.1002/cbic.201500331.

Chen, N., Zhou, M., Dong, X., Qu, J., Gong, F., Han, Y., et al. (2020). Epidemiological and clinical characteristics of 99 cases of 2019 novel coronavirus pneumonia in Wuhan, China: A descriptive study. Lancet 395, 507–513. doi: 10.1016/S0140-6736(20)30211-7.

Chiabrando, D., Vinchi, F., Fiorito, V., Mercurio, S., and Tolosano, E. (2014). Heme in pathophysiology: A matter of scavenging, metabolism and trafficking across cell membranes. Front. Pharmacol. 5, 61. doi: 10.3389/fphar.2014.00061.

Cucinotta, D., and Vanelli, M. (2020). WHO declares COVID-19 a pandemic. Acta Biomed. 91, 157–160. doi: 10.23750/abm.v91i1.9397.

Detzel, M. S., Syllwasschy, B. F., Steinbock, F., Ramoji, A., Hopp, M.-T., Neugebauer, U., et al. (2020). Revisiting the interaction of heme with hemopexin: Recommendations for the responsible use of an emerging drug. bioRxiv. doi: https://doi.org/10.1101/2020.04.16.044321.

Domingo-Fernández, D., Baksi, S., Schultz, B., Gadiya, Y., Karki, R., Raschka, T., et al. (2020). COVID-19 Knowledge Graph: a computable, multi-modal, cause-and-effect knowledge model of COVID-19 pathophysiology. bioRxiv. doi: 10.1101/2020.04.14.040667.

Dutra, F. F., and Bozza, M. T. (2014). Heme on innate immunity and inflammation. Front. Pharmacol. 5, 1–20. doi: 10.3389/fphar.2014.00115.

Fan, B. E., Chong, V. C. L., Chan, S. S. W., Lim, G. H., Lim, K. G. E., Tan, G. B., et al. (2020). Hematologic parameters in patients with COVID-19 infection. Am. J. Hematol. 95, e131–e134. doi: 10.1002/ajh.25774.

Fielding, B. C., Gunalan, V., Tan, T. H. P., Chou, C.-F., Shen, S., Khan, S., et al. (2006). Severe acute respiratory syndrome coronavirus protein 7a interacts with hSGT. Biochem. Biophys. Res. Commun. 343, 1201–1208. doi: 10.1016/j.bbrc.2006.03.091.

Fisher, R. A. (1992). “Statistical methods for research workers,” in Breakthroughs in statistics, 66–70. doi: 10.1002/qj.49708235130.

Giannis, D., Ziogas, I. A., and Gianni, P. (2020). Coagulation disorders in coronavirus infected patients: COVID-19, SARS-CoV-1, MERS-CoV and lessons from the past. J. Clin. Virol. 127, 104362. doi: 10.1016/j.jcv.2020.104362.

Goel, A., Shekhar, S., Singh, O., Garg, S., and Sharma, D. (2018). Hepatitis A virus-induced severe hemolysis complicated by severe glucose-6-phosphate dehydrogenase deficiency. Indian J. Crit. Care Med. 22, 670–673. doi: 10.4103/ijccm.IJCCM_260_18.

Guan, W., Ni, Z., Hu, Y., Liang, W., Ou, C., He, J., et al. (2020). Clinical characteristics of coronavirus disease 2019 in China. N. Engl. J. Med. 382, 1708–1720. doi: 10.1056/NEJMoa2002032.

Gupta, N., de Wispelaere, M., Lecerf, M., Kalia, M., Scheel, T., Vrati, S., et al. (2015). Neutralization of Japanese Encephalitis Virus by heme-induced broadly reactive human monoclonal antibody. Sci. Rep. 5, 16248. doi: 10.1038/srep16248.

Han, H., Yang, L., Liu, R., Liu, F., Wu, K., Li, J., et al. (2020). Prominent changes in blood coagulation of patients with SARS-CoV-2 infection. Clin. Chem. Lab. Med. doi: 10.1515/cclm-2020-0188.

Hänel, K., and Willbold, D. (2007). SARS-CoV accessory protein 7a directly interacts with human LFA-1. Biol. Chem. 388, 1325–1332. doi: 10.1515/BC.2007.157.

Hanff, T. C., Harhay, M. O., Brown, T. S., Cohen, J. B., and Mohareb, A. M. (2020). Is there an association between COVID-19 mortality and the renin-angiotensin system? – A call for epidemiologic investigations. Clin. Infect. Dis. doi: 10.1093/cid/ciaa329.

Heurich, A., Hofmann-Winkler, H., Gierer, S., Liepold, T., Jahn, O., and Pohlmann, S. (2014). TMPRSS2 and ADAM17 cleave ACE2 differentially and only proteolysis by TMPRSS2 augments entry driven by the severe acute respiratory syndrome coronavirus spike protein. J. Virol. 88, 1293–1307. doi: 10.1128/JVI.02202-13.

Hoffmann, M., Kleine-Weber, H., Schroeder, S., Krüger, N., Herrler, T., Erichsen, S., et al. (2020). SARS-CoV-2 cell entry depends on ACE2 and TMPRSS2 and is blocked by a clinically proven protease inhibitor. Cell 181, 271–280.e8. doi: 10.1016/j.cell.2020.02.052.

Hoyt, C. T., Konotopez, A., and Ebeling, C. (2018). PyBEL: A computational framework for Biological Expression Language. Bioinformatics 34, 703–704. doi: 10.1093/bioinformatics/btx660.

Huang, C., Wang, Y., Li, X., Ren, L., Zhao, J., Hu, Y., et al. (2020). Clinical features of patients infected with 2019 novel coronavirus in Wuhan, China. Lancet 395, 497–506. doi: 10.1016/S0140-6736(20)30183-5.

Humayun, F., Domingo-Fernández, D., Paul George, A. A., Hopp, M.-T., Syllwasschy, B. F., Detzel, M. S., et al. (2020). A computational approach for mapping heme biology in the context of hemolytic disorders. Front. Bioeng. Biotechnol. 8, 74. doi: 10.3389/fbioe.2020.00074.

Ji, H.-L., Zhao, R., Matalon, S., and Matthay, M. A. (2020). Elevated plasmin(ogen) as a common risk factor for COVID-19 susceptibility. Physiol. Rev. 100, 1065–1075. doi: 10.1152/physrev.00013.2020.

Krieger, E., and Vriend, G. (2014). YASARA View – molecular graphics for all devices – from smartphones to workstations. Bioinformatics 30, 2981–2982. doi: 10.1093/bioinformatics/btu426.

Kühl, T., and Imhof, D. (2014). Regulatory Fe II/III Heme: The reconstruction of a molecule’s biography. ChemBioChem 15, 2024–2035. doi: 10.1002/cbic.201402218.

Kumar, A., Wißbrock, A., Goradia, N., Bellstedt, P., Ramachandran, R., Imhof, D., et al. (2018). Heme interaction of the intrinsically disordered N-terminal peptide segment of human cystathionine-ß-synthase. Sci. Rep. 8, 1–9. doi: 10.1038/s41598-018-20841-z.

Kupke, T., Klare, J. P., and Brügger, B. (2020). Heme binding of transmembrane signaling proteins undergoing regulated intramembrane proteolysis. Commun. Biol. 3, 73. doi: 10.1038/s42003-020-0800-0.

Kwon, S., Kim, E., Jung, Y. S., Jang, J. S., and Cho, N. (2020). Post-donation COVID-19 identification in blood donors. Vox Sang. doi: 10.1111/vox.12925.

Lan, J., Ge, J., Yu, J., Shan, S., Zhou, H., Fan, S., et al. (2020). Structure of the SARS-CoV-2 spike receptor-binding domain bound to the ACE2 receptor. Nature 581, 215–220. doi: 10.1038/s41586-020-2180-5.

Lecerf, M., Scheel, T., Pashov, A. D., Jarossay, A., Ohayon, D., Planchais, C., et al. (2015). Prevalence and gene characteristics of antibodies with cofactor-induced HIV-1 specificity. J. Biol. Chem. 290, 5203–5213. doi: 10.1074/jbc.M114.618124.

Litalien, C., Proulx, F., Mariscalco, M. M., Robitaille, P., Turgeon, J. P., Orrbine, E., et al. (1999). Circulating inflammatory cytokine levels in hemolytic uremic syndrome. Pediatr. Nephrol. 13, 840–845. doi: 10.1007/s004670050712.

Liu, W., and Li, H. (2020). COVID-19: Attacks the 1-beta chain of hemoglobin and captures the porphyrin to inhibit human heme metabolism. ChemRxiv. doi: 10.26434/chemrxiv.11938173.

Lukassen, S., Chua, R. L., Trefzer, T., Kahn, N. C., Schneider, M. A., Muley, T., et al. (2020). SARS-CoV-2 receptor ACE2 and TMPRSS2 are primarily expressed in bronchial transient secretory cells. EMBO J. 39, e105114. doi: 10.15252/embj.20105114.

Mendes de Oliveira, G., Valle Garay, A., Araújo Souza, A., Cunha, J., Lima, B., Fonseca Valadares, N., et al. (2018). Structural characterization and crystallization of human TMPRSS2 protease. Biophys. J. 114, 567a. doi: 10.1016/j.bpj.2017.11.3102.

Mogl, M. T. (2007). An unhappy triad: Hemochromatosis, porphyria cutanea tarda and hepatocellular carcinoma-A case report. World J. Gastroenterol. 13, 1998–2001. doi: 10.3748/wjg.v13.i13.1998.

Mousavizadeh, L., and Ghasemi, S. (2020). Genotype and phenotype of COVID-19: Their roles in pathogenesis. J. Microbiol. Immunol. Infect. in press. doi: 10.1016/j.jmii.2020.03.022.

Neris, R. L. S., Figueiredo, C. M., Higa, L. M., Araujo, D. F., Carvalho, C. A. M., Verçoza, B. R. F., et al. (2018). Co-protoporphyrin IX and Sn-protoporphyrin IX inactivate Zika, Chikungunya and other arboviruses by targeting the viral envelope. Sci. Rep. 8, 9805. doi: 10.1038/s41598-018-27855-7.

Ou, X., Liu, Y., Lei, X., Li, P., Mi, D., Ren, L., et al. (2020). Characterization of spike glycoprotein of SARS-CoV-2 on virus entry and its immune cross-reactivity with SARS-CoV. Nat. Commun. 11, 1620. doi: 10.1038/s41467-020-15562-9.

Paul George, A. A., Lacerda, M., Syllwasschy, B. F., Hopp, M.-T., Wißbrock, A., and Imhof, D. (2020). HeMoQuest: a webserver for qualitative prediction of transient heme binding to protein motifs. BMC Bioinformatics 21, 124. doi: 10.1186/s12859-020-3420-2.

Ren, X. (2006). Analysis of ACE2 in polarized epithelial cells: surface expression and function as receptor for severe acute respiratory syndrome-associated coronavirus. J. Gen. Virol. 87, 1691–1695. doi: 10.1099/vir.0.81749-0.

Risitano, A. M., Mastellos, D. C., Huber-Lang, M., Yancopoulou, D., Garlanda, C., Ciceri, F., et al. (2020). Complement as a target in COVID-19? Nat. Rev. Immunol. 20, 343–344. doi: 10.1038/s41577-020-0320-7.

Roumenina, L. T., Rayes, J., Lacroix-Desmazes, S., and Dimitrov, J. D. (2016). Heme: Modulator of plasma systems in hemolytic diseases. Trends Mol. Med. 22, 200–213. doi: 10.1016/j.molmed.2016.01.004.

Ruch, T. R., and Machamer, C. E. (2012). The coronavirus E protein: Assembly and beyond. Viruses 4, 363–382. doi: 10.3390/v4030363.

Syllwasschy, B. F., Beck, M. S., Družeta, I., Hopp, M.-T., Ramoji, A., Neugebauer, U., et al. (2020). High-affinity binding and catalytic activity of His/Tyr-based sequences: Extending heme-regulatory motifs beyond CP. Biochim. Biophys. Acta – Gen. Subj. 1864, 129603. doi: 10.1016/j.bbagen.2020.129603.

Tang, N., Li, D., Wang, X., and Sun, Z. (2020). Abnormal coagulation parameters are associated with poor prognosis in patients with novel coronavirus pneumonia. J. Thromb. Haemost. 18, 844–847. doi: 10.1111/jth.14768.

Walls, A. C., Park, Y.-J., Tortorici, M. A., Wall, A., McGuire, A. T., and Veesler, D. (2020). Structure, function, and antigenicity of the SARS-CoV-2 spike glycoprotein. Cell 181, 281–292.e6. doi: 10.1016/j.cell.2020.02.058.

Wang, R., Pan, M., Zhang, X., Han, M., Fan, X., Zhao, F., et al. (2020). Epidemiological and clinical features of 125 Hospitalized Patients with COVID-19 in Fuyang, Anhui, China. Int. J. Infect. Dis. 95, 421–428. doi: 10.1016/j.ijid.2020.03.070.

Waterhouse, A., Bertoni, M., Bienert, S., Studer, G., Tauriello, G., Gumienny, R., et al. (2018). SWISS-MODEL: Homology modelling of protein structures and complexes. Nucleic Acids Res. 46, W296–W303. doi: 10.1093/nar/gky427.

Wißbrock, A., Goradia, N. B., Kumar, A., Paul George, A. A., Kühl, T., Bellstedt, P., et al. (2019). Structural insights into heme binding to IL-36a proinflammatory cytokine. Sci. Rep. 9, 16893. doi: 10.1038/s41598-019-53231-0.

Wu, K., Peng, G., Wilken, M., Geraghty, R. J., and Li, F. (2012). Mechanisms of host receptor adaptation by severe acute respiratory syndrome coronavirus. J. Biol. Chem. 287, 8904–8911. doi: 10.1074/jbc.M111.325803.

Xu, H., Zhong, L., Deng, J., Peng, J., Dan, H., Zeng, X., et al. (2020). High expression of ACE2 receptor of 2019-nCoV on the epithelial cells of oral mucosa. Int. J. Oral Sci. 12, 8. doi: 10.1038/s41368-020-0074-x.

Yan, R., Zhang, Y., Li, Y., Xia, L., Guo, Y., and Zhou, Q. (2020). Structural basis for the recognition of SARS-CoV-2 by full-length human ACE2. Science (80-.). 367, 1444–1448. doi: 10.1126/science.abb2762.

Yang, X., Yu, Y., Xu, J., Shu, H., Xia, J., Liu, H., et al. (2020). Clinical course and outcomes of critically ill patients with SARS-CoV-2 pneumonia in Wuhan, China: a single-centered, retrospective, observational study. Lancet Respir. Med. 8, 475–481. doi: 10.1016/S2213-2600(20)30079-5.

Ye, Q., Wang, B., and Mao, J. (2020). The pathogenesis and treatment of the ‘Cytokine Storm’ in COVID-19. J. Infect. 80, 607–613. doi: 10.1016/j.jinf.2020.03.037.

Young, B. E., Ong, S. W. X., Kalimuddin, S., Low, J. G., Tan, S. Y., Loh, J., et al. (2020). Epidemiologic features and clinical course of patients infected with SARS-CoV-2 in Singapore. J Amer Med Assoc 323, 1488–1494. doi: 10.1001/jama.2020.3204.

Zhang, C., Zheng, W., Huang, X., Bell, E. W., Zhou, X., and Zhang, Y. (2020a). Protein structure and sequence reanalysis of 2019-nCoV genome refutes snakes as its intermediate host and the unique similarity between its spike protein insertions and HIV-1. J. Proteome Res. 19, 1351–1360. doi: 10.1021/acs.jproteome.0c00129.

Zhang, H., Penninger, J. M., Li, Y., Zhong, N., and Slutsky, A. S. (2020b). Angiotensin-converting enzyme 2 (ACE2) as a SARS-CoV-2 receptor: Molecular mechanisms and potential therapeutic target. Intensive Care Med. 46, 586–590. doi: 10.1007/s00134-020-05985-9.

Zhang, X., Cai, H., Hu, J., Lian, J., Gu, J., Zhang, S., et al. (2020c). Epidemiological, clinical characteristics of cases of SARS-CoV-2 infection with abnormal imaging findings. Int. J. Infect. Dis. 94, 81–87. doi: 10.1016/j.ijid.2020.03.040.

Zhou, F., Yu, T., Du, R., Fan, G., Liu, Y., Liu, Z., et al. (2020a). Clinical course and risk factors for mortality of adult inpatients with COVID-19 in Wuhan, China: a retrospective cohort study. Lancet 395, 1054–1062. doi: 10.1016/S0140-6736(20)30566-3.

Zhou, P., Yang, X.-L., Wang, X.-G., Hu, B., Zhang, L., Zhang, W., et al. (2020b). A pneumonia outbreak associated with a new coronavirus of probable bat origin. Nature 579, 270–273. doi: 10.1038/s41586-020-2012-7.

