## Supplementary material for "Unravelling the debate on heme effects in COVID-19 infections"

### 1 Supplementary Tables

**Supplementary Table 1.** Original evidence for the depicted connections for "Immune Response – Inflammation".

| Source | Relation | Target | Original evidence | PMID | Resource |
|---|---|---|---|---|---|
| COVID-19 | Increases | IL-1β | When **COVID-19** infects the upper and lower respiratory tract it can cause mild or highly acute respiratory syndrome with consequent release of pro-inflammatory cytokines, including **interleukin (IL)-1β** and IL-6. | 32171193 | COVID-19 KG |
| Heme | Increases | IL-1β | **Heme** activates macrophages inducing the production of TNF, KC (Figueiredo et al., 2007), **IL-1β** (unpublished), and LTB4. | 24904418 | Heme KG |
| COVID-19 | Increases | IL-6 | When **COVID-19** infects the upper and lower respiratory tract it can cause mild or highly acute respiratory syndrome with consequent release of pro-inflammatory cytokines, including interleukin (IL)-1β and **IL-6.** | 32171193 | COVID-19 KG |
| Heme | Increases | IL-6 | Likewise, **heme** and FeNTA treatment causes the induction of the M1 | 26675351 | Heme KG |

| | | | | | |
|---|---|---|---|---|---|
| | | | markers MHC II, CD86, CD14, TNFα, **IL-6**, and IL1β and a decrease in the M2 markers CD206, IL-10, and Arginase-1 (the last with FeNTA only) in M0 BMDMs (Figure 3A; supplemental Figures 5, 6A, and 7). | | |
| COVID-19 | Increases | TNFα | Initial plasma IL1B, IL1RA, IL7, IL8, IL9, IL10, basic FGF, GCSF, GMCSF, IFNγ, IP10, MCP1, MIP1A, MIP1B, PDGF, **TNFα**, and VEGF concentrations were higher in both ICU patients and non-ICU patients than in healthy adults (appendix pp 6–7). | 31986264 | COVID-19 KG |
| Heme | Increases | TNFα | On the other hand, **TNF** secretion induced by **heme** is essential for the activation of the programmed necrotic cell death pathway, which is denominated necroptosis.<br><br>Likewise, **heme** and FeNTA treatment causes the induction of the M1 markers MHC-II, CD86, CD14, **TNFα**, IL-6, and IL1β and a decrease in the M2 markers CD206, IL-10, and Arginase-1 (the last with FeNTA only) in M0 BMDMs (Figure 3A; supplemental Figures 5, 6A, and 7). | 24904418<br><br>26675351 | Heme KG<br><br>Heme KG |

| | | | | | |
|---|---|---|---|---|---|
| TNFα | Increases | TLR4 | Heme also induces tumour necrosis factor (**TNF**) secretion in monocyte/macrophages through **TLR4** and the adaptor molecule, MYD88/. | 25307023 | Heme KG |
| Heme | Increases | TLR4 | **Heme** also induces tumour necrosis factor (TNF) secretion in monocyte/macrophages through **TLR4** and the adaptor molecule, MYD88/. | 25307023 | Heme KG |
| TLR4 | Increases | MyD88 | The **TLR4** activates two distinct pathways: **MyD88** and TRIF. In macrophages, heme induces a biased MyD88 activation and the secretion of the pro-inflammatory cytokines TNF and KC. | 24904418 | Heme KG |
| MyD88 | Increases | NF-κB | Only under protein-free conditions did we observe a limited heme-induced TNF-alpha response in cultured macrophages, which was triggered via signaling of the classical TLR4–**MyD88**–TRIF pathway of **NF-kB** activation. | 29610666 | Heme KG |
| NF-κB | Increases | IL-6 | The MyD88-dependent pathway leads to the activation of mitogen-activated protein kinases (MAPKs) and **NF-κB** (nuclear factor kappa- | 24904418 | Heme KG |

| | | | | | |
|---|---|---|---|---|---|
| | | | light-chain-enhancer of activated B cells) transcription factors inducing the expression of inflammatory cytokines such as TNF, **IL-6**, IL-1β, and KC | | |
| COVID-19 | Increases | IL-8 | Initial plasma IL1B, IL1RA, IL7, **IL8**, IL9, IL10, basic FGF, GCSF, GMCSF, IFNγ, IP10, MCP1, MIP1A, MIP1B, PDGF, TNFα, and VEGF concentrations were higher in both ICU patients and non-ICU patients than in healthy adults (appendix pp 6–7). | 31986264 | COVID-19 KG |
| Heme | Increases | IL-8 | Incubation of neutrophils with 4 mM **hemin** resulted in a 2-fold increase in **CXCL8** mRNA expression and in a significant increase in released and cell-associated levels of IL-8 protein compared with the untreated controls or A1AT-treated cells (Fig. 7A–C). | 28716864 | Heme KG |
| COVID-19 | Increases | CD14 | Although patients had higher **sCD14** levels than healthy people, there were no significant differences between the severe and mild groups (Fig. 1b). | 32203186 | COVID-19 KG |
| Heme | Increases | CD14 | Likewise, **heme** and FeNTA treatment causes | 26675351 | Heme KG |

| | | | | | |
|---|---|---|---|---|---|
| | | | the induction of the M1 markers MHC-II, CD86, **CD14**, TNFα, IL-6, and IL1β and a decrease in the M2 markers CD206, IL-10, and Arginase-1 (the last with FeNTA only) in M0 BMDMs (Figure 3A; supplemental Figures 5, 6A, and 7). | | |
| COVID-19 | Increases | IL-10 | Significantly high blood levels of cytokines and chemokines were noted in patients with **COVID-19** infection that included IL1-β, IL1RA, IL7, IL8, IL9, **IL10**, basic FGF2, GCSF, GMCSF, IFNγ, IP10, MCP1, MIP1α, MIP1β, PDGFB, TNFα, and VEGFA. Some of the severe cases that were admitted to the intensive care unit showed high levels of pro-inflammatory cytokines including IL2, IL7, **IL10**, GCSF, IP10, MCP1, MIP1α, and TNFα that are reasoned to promote disease severity.<br><br>Initial plasma IL1B, IL1RA, IL7, IL8, IL9, **IL10**, basic FGF, GCSF, GMCSF, IFNγ, IP10, MCP1, MIP1A, MIP1B, PDGF, TNFα, and VEGF concentrations were higher in both ICU patients and non-ICU patients than in healthy adults (appendix pp 6–7). | 32113704<br><br>31986264 | COVID-19 KG<br><br>COVID-19 KG |

| | | | | | |
|---|---|---|---|---|---|
| Heme | Increases | IL-10 | Interestingly, **heme** (10–30 μM) was also able to induce the secretion of **IL-10**, an anti-inflammatory cytokine. | 23690673 | Heme KG |

**Supplementary Table 2.** Original evidence for the depicted connections for "Immune Response – Complement system".

| **Source** | **Relation** | **Target** | **Original Evidence** | **PMID/doi** | **Resource** |
|---|---|---|---|---|---|
| COVID-19 | Increases | C3 | **C3** activation products (C3 fragments C3a, C3b, iC3b, C3dg, and C3c) were detected by Western blotting in lung tissue of SARS-CoV MA15-infected mice, but not in control mice, as early as 1 day post infection (dpi) (Fig. 1B), confirming that SARS-CoV MA15 infection activates the complement pathway. | 30301856 | COVID-19 KG |
| Heme | Increases | C3 | Furthermore, the incubation of serum or whole blood with **heme** induces deposition of activation fragments (C3b, iC3b, C3dg) of **complement component 3** (C3) at the surface of erythrocytes | 26875449 | Heme KG |
| COVID-19 | Increases | Comple-ment activation | C3 activation products (C3 fragments C3a, C3b, iC3b, C3dg, and C3c) were detected by Western blotting in lung tissue of | 30301856 | COVID-19 KG |

| | | | | | |
|---|---|---|---|---|---|
| | | | SARS-CoV MA15-infected mice, but not in control mice, as early as 1 day post infection (dpi) (Fig. 1B), confirming that SARS-CoV MA15 infection **activates the complement pathway**. | | |
| Heme | Increases | Comple-ment activation | Hemolytic diseases are often accompanied by dysregulation and **overactivation of the complement system** [72–74], which may be induced by free extracellular **heme**. | 26875449 | Heme KG |
| COVID-19 | Increases | Neutrophil activation | The common clinical manifestations of fever and cough in conjunction with laboratory results of progressively increasing neutrophil counts and leukocytopenia in serum samples taken from **COVID-19** patients at various stages of the illness indicates uncontrolled **neutrophil** extravasation and **activation**. | 10.1101/2020.03.31.019216 | COVID-19 KG |
| Heme | Increase | Neutrophil activation | **Heme-induced neutrophils activation** leads to extracellular traps (NETs) release through a mechanism dependent on ROS generation. | 24904418 | Heme KG |
| Heme | Increases | C5a | C3a and **C5a** anaphylatoxins, as well as the soluble membrane attack complex (sC5b9), | 26875449 | Heme KG |

| | | | |
|---|---|---|---|
| | | | are generated by incubation of **heme** with human serum or blood in vitro, via the alternative complement pathway). |

**Supplementary Table 3.** Original evidence for the depicted connections for "Blood and coagulation system".

| **Source** | **Relation** | **Target** | **Original Evidence** | **PMID** | **Resource** |
|---|---|---|---|---|---|
| COVID-19 | Decreases | Albumin | **Albumin** concentrations were significantly lower in deceased patients than in recovered patients. | 32217556 | COVID-19 KG |
| Albumin | Decreases | Heme | Serum **albumin** (SA) can act as the heme scavenger by<br><br>forming **heme**-SA complex [2, 4–8]. | 30324533 | Heme KG |
| COVID-19 | Increases | Ferritin | Levels of lactate dehydrogenase (LDH), concentrations of serum high-sensitivity C-reactive protein (hsCRP), **ferritin** and D-dimer levels were markedly higher in severe cases than moderate cases. | 32217835 | COVID-19 KG |
| Heme | Increases | Ferritin | In agreement with the electrical cell–substrate impedance sensing data described above, the proteome changes triggered by 10 μM **heme** were indicative of an | 26794659 | Heme KG |

| | | | | | |
|---|---|---|---|---|---|
| | | | adaptive response with prominent induction of HMOX1 and **ferritin** light (FTL) and heavy (FTH1) chains. | | |
| COVID-19 | Decreases | Plasminogen activation | Figure 2. GO-term and KEGG pathway enrichment of up-regulated expressed genes in BALF and PBMC of COVID-19 patients. [figure content] | 32228226 | COVID-19 KG |
| COVID-19 | Decreases | Platelet count | Remarkable abnormalities in the CBC were detected on January 29, including increased neutrophils ($10.67\times10^9$/L), leukocytes ($11.73\times10^9$/L), and decreased lymphocytes ($0.51\times10^9$ /L), and on February 4 decreased erythrocytes ($2.40\times10^{12}$/L) and **platelets** ($73\times10^9$ /L). | 32196678 | COVID-19 KG |
| Heme | Increases | Platelet aggregation | 2 **Heme** induces **platelet** activation and **aggregation** through different pathways | 26875449 | Heme KG |
| Heme | Decreases | Fibrin | From another perspective, in vitro assays have demonstrated that **heme** can also bind to fibrinogen and decrease its thrombin-mediated cleavage, thus affecting the final common coagulation pathway and reducing **fibrin formation**, important in clotting (Figure 1). | 26875449 | Heme KG |

| | | | | | |
|---|---|---|---|---|---|
| Fibrin | Increases | Hemoglobin | During DIC, **fibrin** strands within the fibrin mesh formed could cut red blood cells, resulting in the formation of schistocytes (strongly deformed red blood cells or fragments of red blood cells) and the release of **hemoglobin**. | 29956069 | Heme KG |
| Hemoglobin | Increases | Heme | Toxicity of free **hemoglobin** is also caused by the release of cell-free **heme**, which produces lipid peroxidation and mitochondrial damage and increases the production of reactive oxygen species. | 27515135 | Heme KG |
| Hemoglobin | Increases | Platelet aggregation | When RBCs are damaged by high shear in continuous flow ventricular assist devices, free **hemoglobin** induces **platelet aggregation**, contributing to high risk of thrombotic complications. | 28458720 | Heme KG |

**Supplementary Table 4.** Original evidence for the depicted connections for "Organ-specific diagnostic markers".

| **Source** | **Relation** | **Target** | **Original Evidence** | **PMID** | **Resource** |
|---|---|---|---|---|---|
| Hemolysis | Increases | Heme | Free **heme** is generated by intra- and extravascular **hemolysis** or extensive cell damage. | 24464629 | Heme KG |

| | | | | | |
|---|---|---|---|---|---|
| Hemolysis | Increases | Bilirubin | LDH was strongly associated with markers normally elevated in either **hemolysis** or liver disease, including aspartate aminotransferase (AST) and direct and indirect **bilirubin**. | 16291595 | Heme KG |
| COVID-19 | Increases | Bilirubin | Concentrations of alanine aminotransferase, aspartate aminotransferase, total **bilirubin**, alkaline phosphatase, and γ-glutamyl transpeptidase were markedly higher in deceased patients than in recovered patients. | 32217556 | COVID-19 KG |
| Heme | Increases | Bilirubin | The present findings demonstrate that the enzymatic mechanism catalyzing the conversion of **heme** to **bilirubin** in the liver is under hormonal control. | 4334719 | Heme KG |
| Hemolysis | Increases | LDH | **LDH** was strongly associated with markers normally elevated in either **hemolysis** or liver disease, including aspartate aminotransferase (AST) and direct and indirect bilirubin. | 16291595 | Heme KG |
| COVID-19 | Increases | LDH | Levels of lactate dehydrogenase (**LDH**), concentrations of serum high-sensitivity C-reactive protein (hsCRP), ferritin and D-dimer levels were | 32217835 | COVID-19 KG |

| | | | |
|---|---|---|---|
| | | | markedly higher in severe cases than moderate cases. |

**Supplementary Table 5.** Concordance between the common interactions and experimental data from Blanco-Melo et al. 2020 (1).

| Pathway | Gene symbol | Cell MMC2 | Consistency MMC2 | Cell MMC4 | Consistency MMC4 |
|---|---|---|---|---|---|
| Immune system - Inflammation | NF-κB | 0.4984 | yes | 0.6478 | yes |
| | MyD88 | 0.199 | yes | 1.519 | yes |
| | IL-6 | 2.390 | yes | -0.517 | no |
| | TLR4 | 0.1533 | yes | 1.917 | yes |
| | IL-1β | 1.036 | yes | 2.354 | yes |
| | CD14 | -0.4775 | no | 0.297 | yes |
| | IL-10 | 0.539 | yes | 4.038 | yes |
| | TNFα | 1.333 | yes | 4.005 | yes |
| Immune system - Complement system | C3 | 1.455 | yes | 0.760 | yes |
| | C5 | 0.050 | -- | -1.392 | -- |
| Blood coagulation | Hemoglobin | -- | -- | -- | -- |

| | | | | | |
|---|---|---|---|---|---|
| system | Ferritin | 0.047 | yes | 1.690 | yes |
| | Albumin | 0.0363 | no | -0.1999 | yes |
| Organ-specific diagnostic markers | LDHA | 0.225 | yes | 0.658 | yes |
| | LDHB | 0.0 | -- | -0.738 | no |
| | LDHC | 0.043 | yes | -0.979 | no |

#### 2 Supplementary References

1. Blanco-Melo D, Nilsson-Payant BE, Liu W-C, Uhl S, Hoagland D, Møller R, Jordan TX, Oishi K, Panis M, Sachs D, et al. Imbalanced host response to SARS-CoV-2 drives development of COVID-19. *Cell* (2020) **181**: 1036-1045.e9. doi:10.1016/j.cell.2020.04.026
